# Static All-Atom Energetic Mappings of the SARS-Cov-2 Spike Protein with Potential Latch Identification of the Down State Protomer

**DOI:** 10.1101/2020.05.12.091090

**Authors:** Michael H. Peters, Oscar Bastidas, Daniel S. Kokron, Christopher Henze

**Affiliations:** *Virginia Commonwealth University*, Department of Chemical and Life Science Engineering, 601 West Main St, Richmond, Virginia, United States of America; *University of Minnesota*, College of Biological Sciences, University of Minnesota, Minneapolis, Minnesota, United States of America; NASA Ames Research Center, Moffett Field, California, United States of America

**Keywords:** SARS-Cov-2, spike protein, energetic mappings, contact plot, latch, COVID-19

## Abstract

The SARS-Cov-2 virion responsible for the current world-wide pandemic Covid-19 has a characteristic Spike protein (S) on its surface that embellishes both a prefusion state and fusion state. The prefusion Spike protein (S) is a large trimeric protein where each protomer may be in a so-called Up state or Down state, depending on the configuration of its receptor binding domain (RBD). The Up state is believed to allow binding of the virion to ACE-2 receptors on human epithelial cells, whereas the Down state is believed to be relatively inactive or reduced in its binding behavior. We have performed detailed all-atom, dominant energy landscape mappings for noncovalent interactions (charge, partial charge, and van der Waals) of the SARS-Cov-2 Spike protein in its static prefusion state based on recent structural information. We included both interchain interactions and intrachain (domain) interactions in our mappings in order to determine any telling differences (different so-called “glue” points) between residues in the Up and Down state protomers. In general, the S2 or fusion machinery domain of S is relatively rigid with strong noncovalent interactions facilitated by helical secondary structures, whereas the S1 domain, which contains the RBD and N-terminal domain (NTD), is relatively more flexible and characterized by beta strand structural motifs. The S2 domain demonstrated no appreciable energetic differences between Up and Down protomers, including interchain as well as each protomer’s intrachain, S1-S2 interactions. However, the S1 domain interactions across neighboring protomers, which include the RBD-NTD cross chain interactions, showed significant energetic differences between Up-Down and Down-Down neighboring protomers. Surprisingly, the Up-Down, RBD-NTD interactions were overall stronger and more numerous than the Down-Down cross chain interactions, including the appearance of the three residue sequence ALA520-PRO521-ALA522 associated with a turn structure in the RBD of the Up state protomer. Additionally, our *intra*chain dominant energy mappings within each protomer, identified a significant “glue” point or possible “latch” for the Down state protomer between the S1 subdomain, SD1, and the RBD domain of the same protomer that was completely missing in the Up state protomer analysis. Ironically, this dominant energetic interaction in the Down state protomer involved the backbone atoms of the same three residue sequence ALA520-PRO521-ALA522 of the RBD with the R-group of GLN564 in the SD1 domain. Thus, this same three residue sequence acts as a stabilizer of the RBD in the Up conformation through its interactions with its neighboring NTD chain and a kind of latch in the Down state conformation through its interactions with its own SD1 domain. The dominant interaction energy residues identified here are also conserved across reported variations of SARS-Cov-2, as well as the closely related virions SARS-Cov and the bat corona virus RatG13. To help verify the potential latch for the Down state protomer, we conducted some preliminary molecular dynamic simulations that effectively turn off this specific latch glue point via a single point mutation of GLN564. Interestingly, the single point mutation lead to the latch releasing in less than a few nanoseconds, but the latch remained fixed in the wild state protomer for up to 0.1 microseconds that were simulated. Many more detailed studies are needed to understand the dynamics of the Up and Down states of the Spike protein, including the stabilizing chain-chain interactions and the mechanisms of transition from Down to Up state protomers. Nonetheless, static dominant energy landscape mappings and preliminary molecular dynamic studies given here may represent a useful starting point for more detailed dynamic analyses and hopefully an improved understanding of the structure-function relationship of this highly complex protein associated with COVID-19.

## 1. Introduction

SARS-Cov-2 is an RNA virus responsible for the disease Covid-19 which was recently declared a world-wide pandemic by the WHO. SARS-Cov-2 virion enters human epithelial cells by attachment of it’s prefusion-form Spike protein (S) with a specific cell surface receptor ACE2 (Angiotensin Converting Enzyme 2) (1-3). The Spike protein is a large, trimeric protein whose Receptor Binding Domain (RBD) undergoes somewhat unusual dynamic transformations sometimes called “breathing”. From a protein engineering perspective, so-called “breathing” reflects the inherent flexibility and/or localized mobility associated with the Receptor Binding Domain (RBD) of the Spike Protein. In the so-called “Up-state” of the RBD, the (prefusion) protein is able to bind to ACE2 (Angiotensin Converting Enzyme 2) and infect (via a transformation to its fusion state) human epithelial cells (Type I and II pneumocytes; also, alveolar macrophage and nasal mucosal cells), but in the “Down-state” the Spike protein is believed to be inactive to ACE2 binding and to cellular infection. We note that the S1 domain of the Spike protein is shed in the transition from the prefusion state to the fusion state of this virion; those transformational aspects are not considered here.

The exact mechanism and specific structural details associated with the flexibility or local mobility of the RBD in the Up and Down states in SARS-Cov-2 remain unanswered. For example, it is not known whether these states exist simply randomly or by deterministic changes orchestrated by the virion or its environment. Recently unpublished long time Molecular Dynamics (MD) studies (10*µs*) of an isolated Spike Protein by the Shaw Group (4) noted that the protomers tended to persist in their initial states, i.e, Down states remain Down and Up states remain Up. However, the Up state protomer demonstrated further distal displacement and mobility from its initial state that was given by experimental structural data (4).

In order to better understand the differences between the Up and Down protomer states, we conducted an all-atom interacting energy landscape mapping of the entire Spike protein from its *.pdb (Protein Data Bank) structure file (6vsb.pdb) in order to identify interaction energy “glue” points associated with relatively strong non-covalent atom-atom interactions between residues, which may be responsible for specific persistent domains of this complex trimeric protein. In doing so, we were able to identify some unique and potentially critical differences between the Up and Down protomers within the overall trimeric structure, including a possible molecular latch that helps to maintain the RBD in the down state conformation. The latch residues are conserved across the closely related virions SARS-Cov-1 and the bat corona virus RatG13, as well as known variations of the novel corona virus. Comparative analyses between Up and Down state protomers, such as those given here, may lead to potentially new therapeutic targets aimed at disrupting the viral functionality of the Spike protein to its role in COVID-19.

## 2. Materials and Methods

Following our recently published study on *Aβ*42 amyloid fibrils (5), we analyzed the recently published trimeric structure of SARS-Cov-2 (PDB ID: 6vsb) according to the Coulombic (charge and partial atomic charge) and Lennard-Jones (Born and van der Waals forces) atom-atom interaction forces as laid out in the open-source energy mapping algorithm developed by Krall et al (6). This mapping algorithm efficiently parses the strongest non-covalent atom-atom interactions and their inter-atomic distances from structure file data according to empirically established criteria based on the *AMBER*03 force field model to ensure that all dominant interactions are accounted for. These energetic mappings, for example, allowed us to discern important structural differences leading to greater adhesive strengths of *Aβ*42 oligomers versus their *Aβ*40 counterparts. Following our previous studies, the parsing criteria were taken as the upper limit of − 0.1*kT* units for Lennard-Jones (van der Waals) criteria and − 0.3*kT* units for Coulombic interactions, although lower values can also be specified in the analysis part of the mappings in order to further refine the results (6).

The SARS-Cov-2 Spike protein structure consists of three chains or protomers (A, B, and C chains) of which the chain A is given in the so-called “Up” state of its RBD (6vsb.pdb), and chains B and C are in their “Down” state. We energetically mapped the interchain interactions “Up-Down” and “Down-Down” and specific domain interactions (*intra*chain interactions) for the Up and Down state protomers, including S1 and S2 domain interactions and sub domains of S1 that include the RBD domain. In addition, following our static analysis, we conducted some preliminary molecular dynamics studies on a potential “latch” for the Down state protomer. Explicit solvent molecular dynamics (MD) simulations of novel coronavirus spike protein were performed using the NAMD2 program (8). We used the CHARMM-Gui (9) with the CHARMM36m force field along with TIP3P water molecules to explicitly solvate the proteins and add any missing residues from the experimental structure files. Simulations were carried out maintaining the number of simulated particles, pressure and temperature (the NPT ensemble) constant with the Langevin piston method specifically used to maintain a constant pressure of 1 atm. We employed periodic boundary conditions for a water box simulation volume as well as the particle mesh Ewald (PME) method with a 20 Å cutoff distance between the simulated protein and water box edge. The integration time step was 2 femtoseconds with our protein simulations conducted under physiological conditions (37 C, pH of 7.4, physiological ionic strength).

## 3. Results

### Interchain Interactions

Mappings of the dominant atom-atom interactions among the residues for the Up-Down (A-B) and Down-Down (B-C) states are illustrated in Figs. 1 and 2, respectively (Table Suppl.1 A-B and Table Suppl.2 B-C). As expected, the majority of the dominant interactions are within the S2 domain, and the entire three chain structure is greatly stabilized by this feature. There are a number of stabilizing interactions involving the SD1/SD2 domain of any protomer with the S2 and NTD domain of neighboring protomers (Supplementary Figs. Suppl.1 and Suppl.2). The pattern of the S1-S1 interchain interactions involving the RBD of any chain to the N-terminal domain (NTD) of its adjacent neighbor is potentially important as shown in Figs. 1 and 2 for clarity. We specifically examined the interchain interactions between the RBD with the NTD of its neighbor protomer for both the Up-Down (A-B) and Down-Down (B-C) interactions, shown in Figs. 3 and 4, respectively. Somewhat surprisingly, the Up-Down interactions were stronger and more numerous overall than the Down-Down interactions of these close neighbor domains. Additionally, the energetically dominant Up-Down RBD-NTD interactions involve three additional residues (ALA520-PRO521-ALA522) not seen in the Down-Down cross chain interactions. These three residues, however, will be shown below to play a stabilizing *intra*chain role for the Down state protomer.

**Figure 1:**
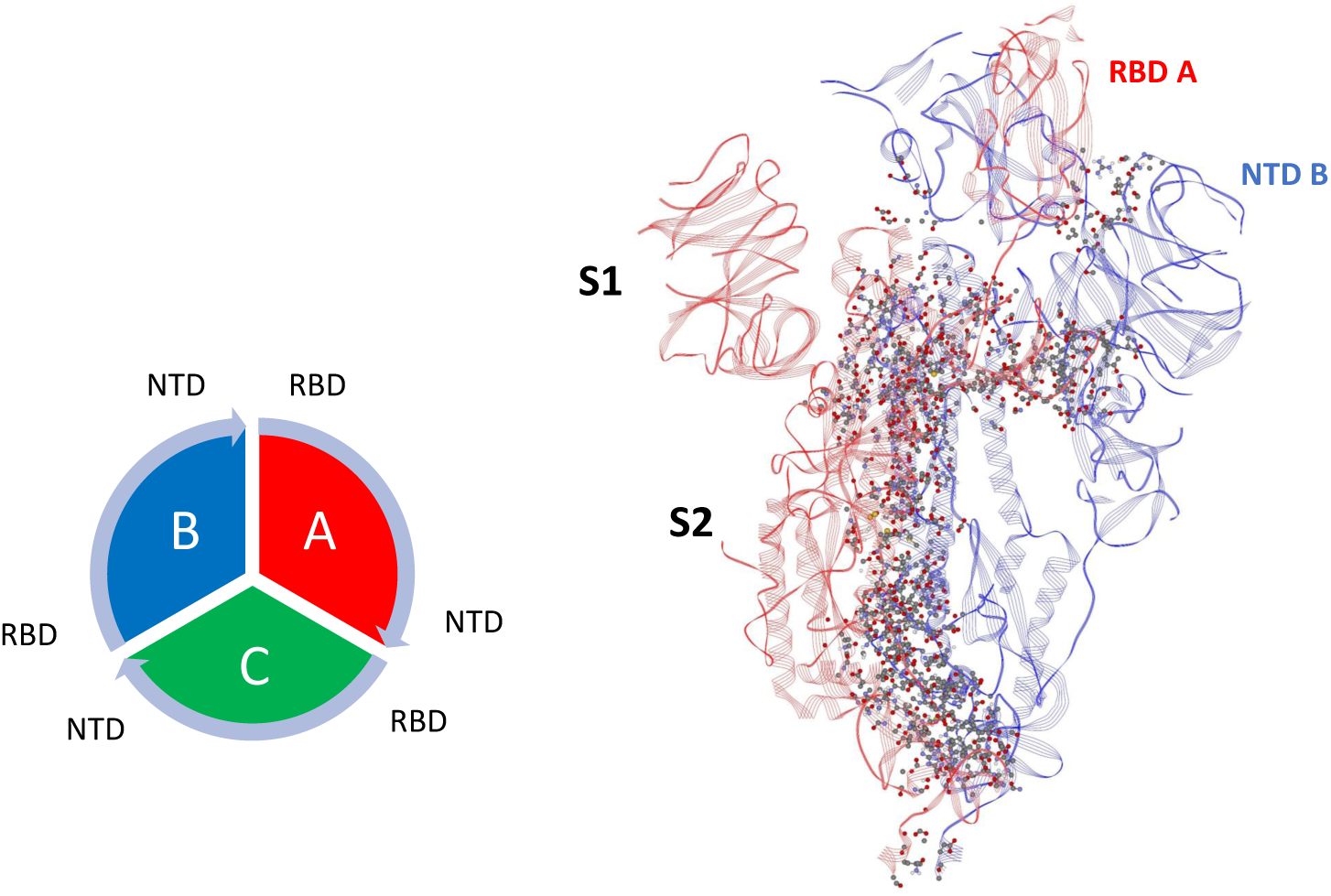
Dominant Energy Landscape for the Interaction of Chain A (Up; red) with Chain B (Down; blue). The dominant atom-atom interactions are shown by ball and sticks. Also shown is the overall chain interaction configuration looking at the trimer from the top view.

**Figure 2:**
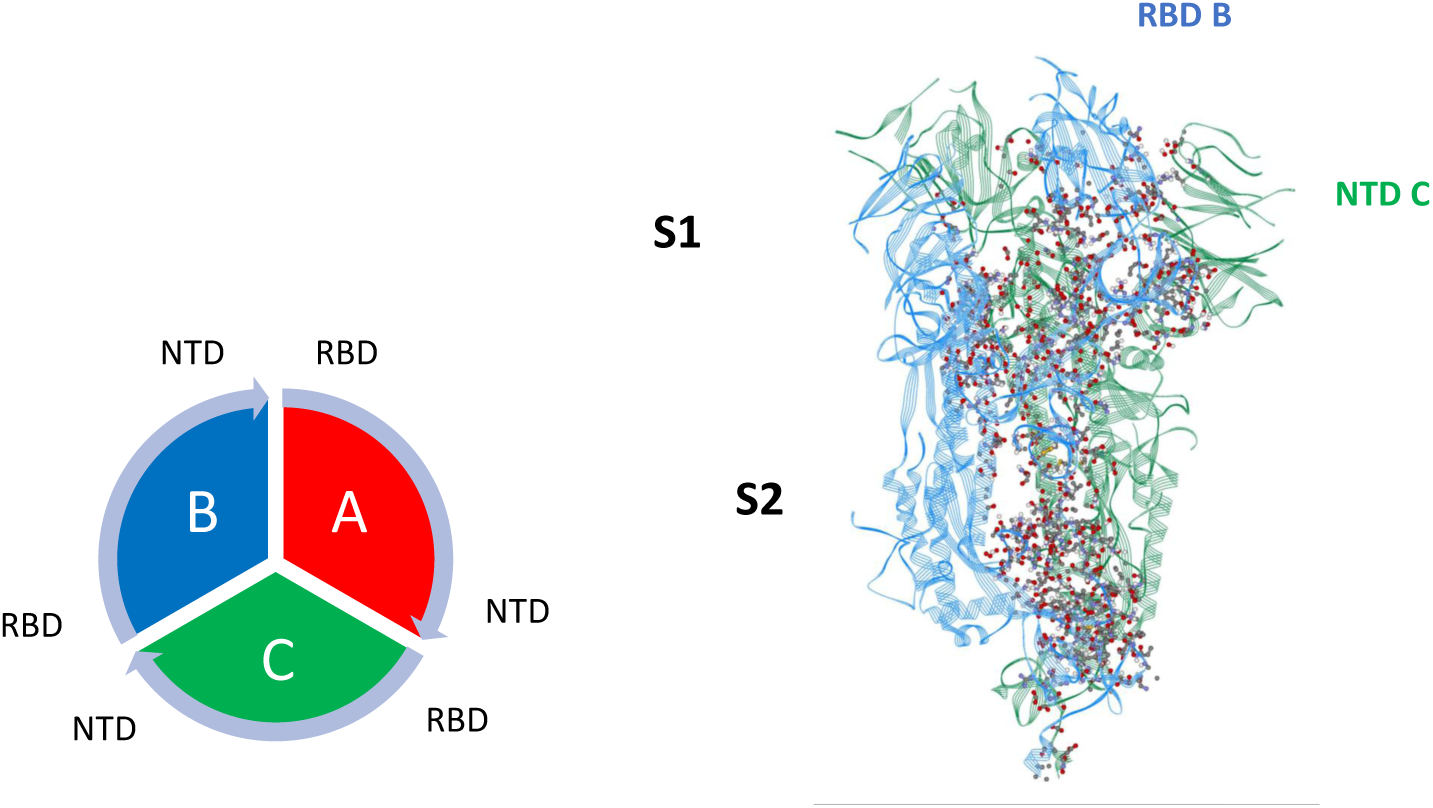
Dominant Energy Landscape for the Interaction of Chain B (Down; blue) with Chain C (Down; green). The dominant atom-atom interactions are shown by ball and sticks.

**Figure 3:**
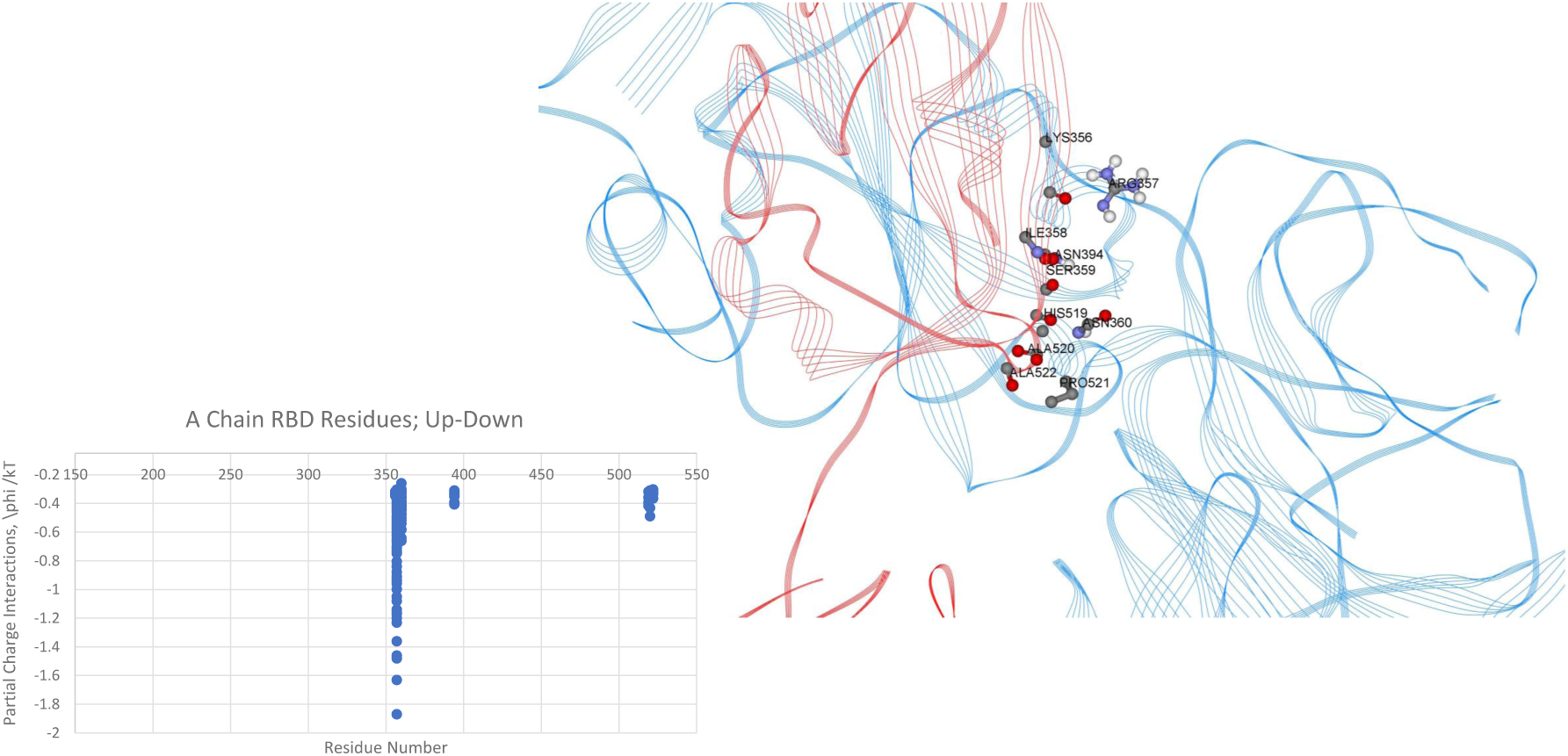
Dominant Energy Landscape for the Interaction of the RBD Chain A (Up;red) with NTD Chain B (Down;blue); oblique, top view. Only A Chain RBD residue/atoms are shown.

**Figure 4:**
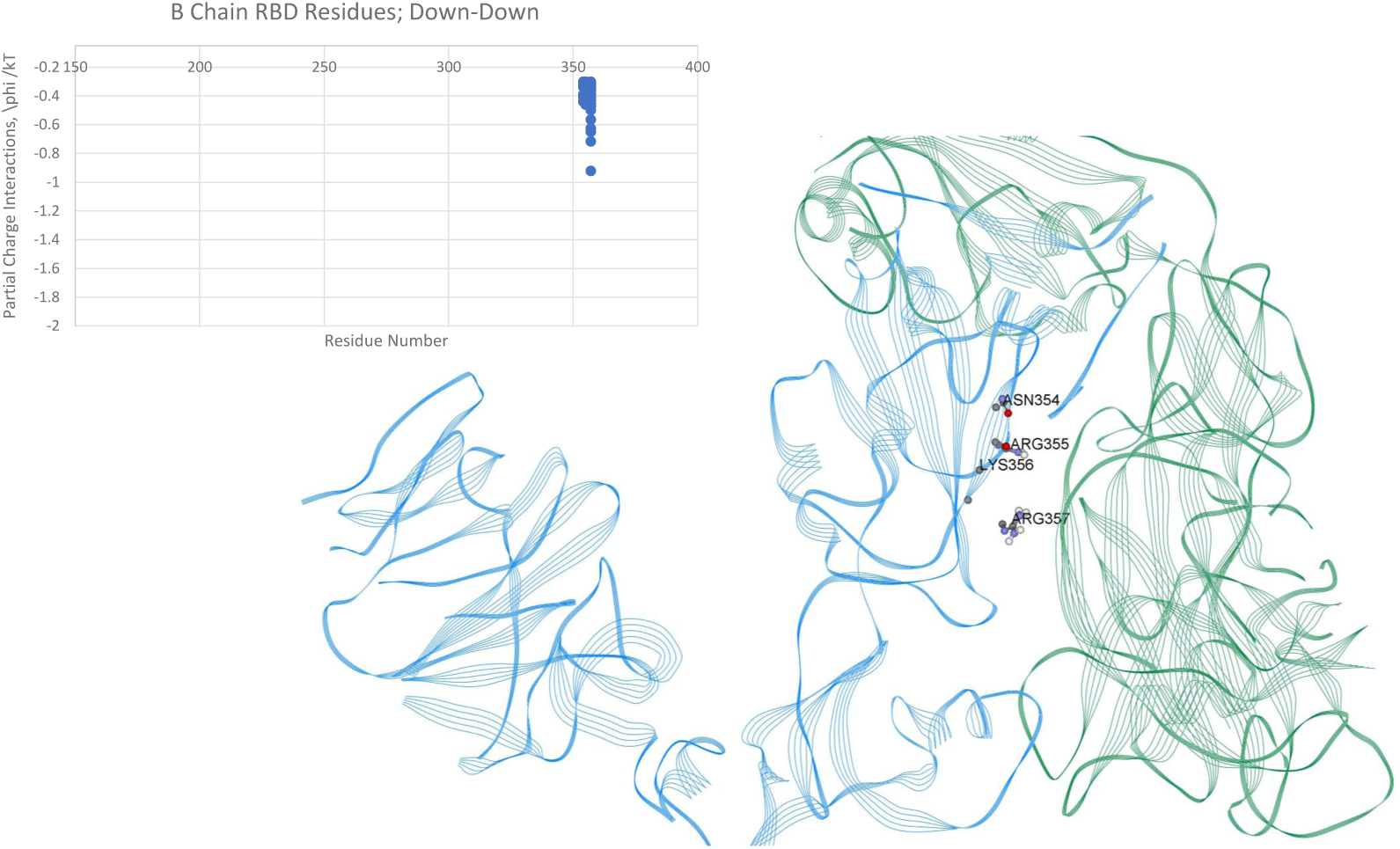
Dominant Energy Landscape for the Interaction of the RBD of Chain B (Down;blue) with the NTD of Chain C (Down;green); oblique, top view. Only B Chain residue/atoms are shown.

### Intrachain Interactions

We next looked in detail at *intra*chain interactions for both the Up and Down protomers in order to attempt to discern any telling differences in these two states.

### S1-S2 Intrachain Interactions

We first energetically mapped the S1 domain (residues 1-677; not including missing residues) to the S2 domain (residues 682-1273; not including missing residues) of both the Up protomer and Down protomer separately, as shown in Tables Suppl.3 and Suppl.4, respectively. Overall, we found no discernable differences in the energetic interactions between the Up and Down states, which again may not be surprising considering the RBD of S1 (residues 319-527) is distal to the S2 domain whether in the Up or Down protomer state. Additionally, the proximal S1 domain beyond the RBD, SD1/SD2 domains (residues 533-677), also showed no discernable differences between the Up and Down states in it’s interaction with S2. We, therefore, next compared the SD1/SD2 domains to RBD domain interactions within each protomer in the Up or Down state, since these states have clear differences in separation distances owing to the more distal RBD in the Up state.

### SD1/SD2-RBD Intrachain Interactions

Overall the energetic mappings between Up and Down states for these two S1 domains (RBD-SD1/SD2) appear very similar, as shown, for example, by the dominant partial charge interactions (Figs. 5 and 6; Tables Suppl.5 and Suppl.6) with one important difference as shown in the zoom illustration of Fig 7.

**Figure 5:**
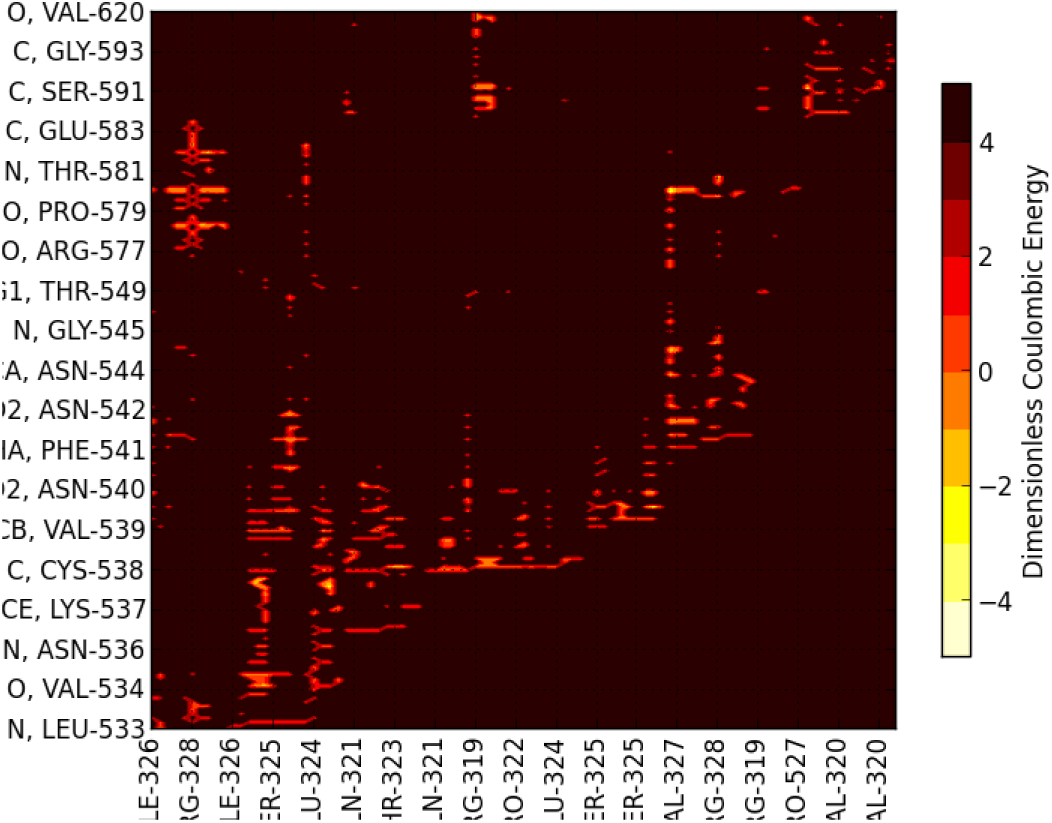
Dominant Energy Landscape for partial charge interactions between SD1/SD2 and RBD of Chain A (Up).

**Figure 6:**
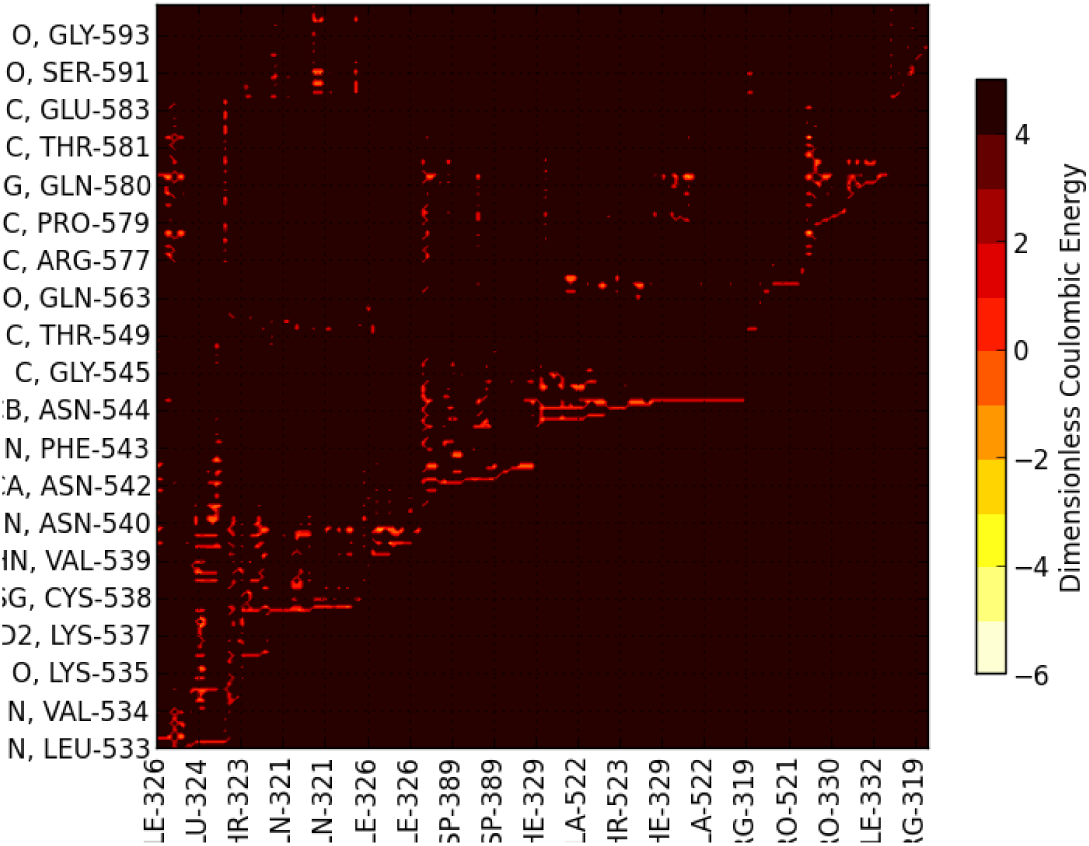
Dominant Energy Landscape for partial charge interactions between SD1/SD2 and RBD of Chain B (Down).

**Figure 7:**
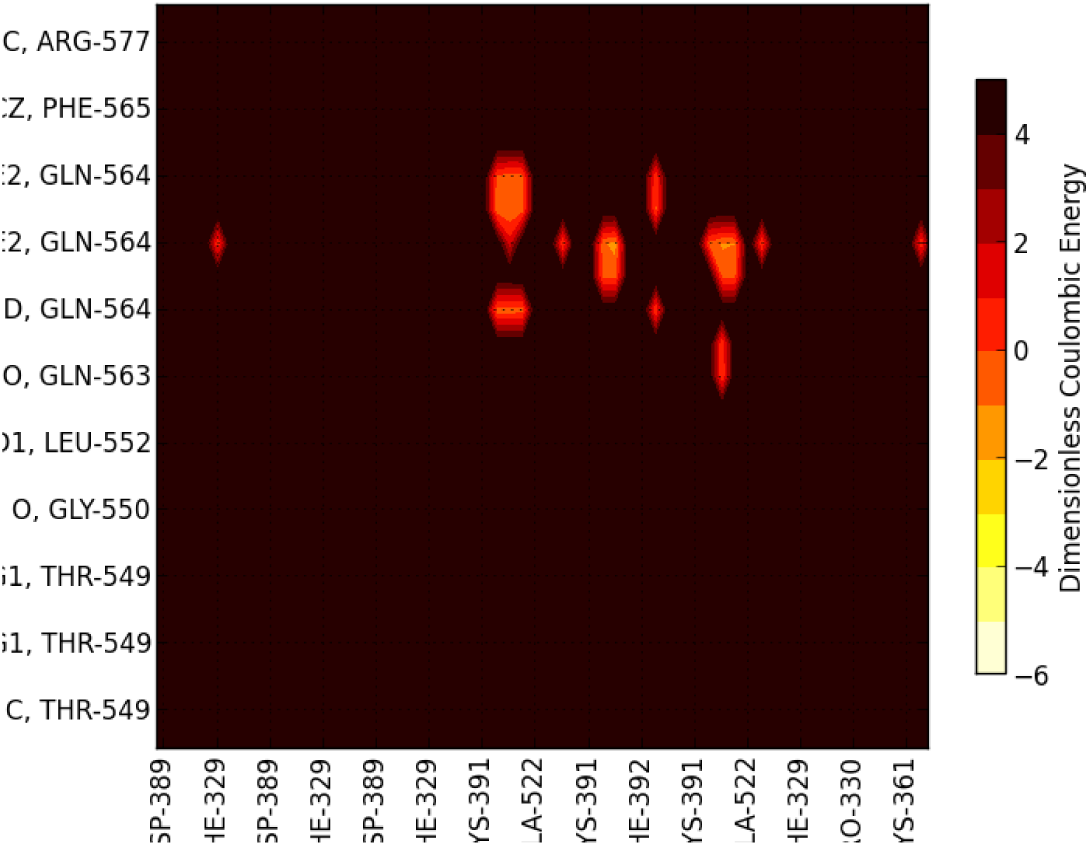
Zoomed Dominant Energy Landscape for partial charge interactions between SD1/SD2 and RBD of Chain B showing the conspicuously unique residue interactions. Note that dominant van der Waals interactions invloving PRO 521 backbone atoms are not shown here; see Tables Suppl.5 and Suppl.6.

For the Down state protomer, we observe conspicuous, strong energetic interactions between the R group of GLN564 in the SD1 domain with backbone atoms of the three sequential residues ALA520-PRO521-ALA522 in the RBD domain, which are missing in the Up state mappings due to the more distal state of the RBD. These specific atom-atom interactions are shown in more detail in Figs. 8 and 9. Note the PRO521 interactions are dominated by van der Waals interactions in both instances.

**Figure 8:**
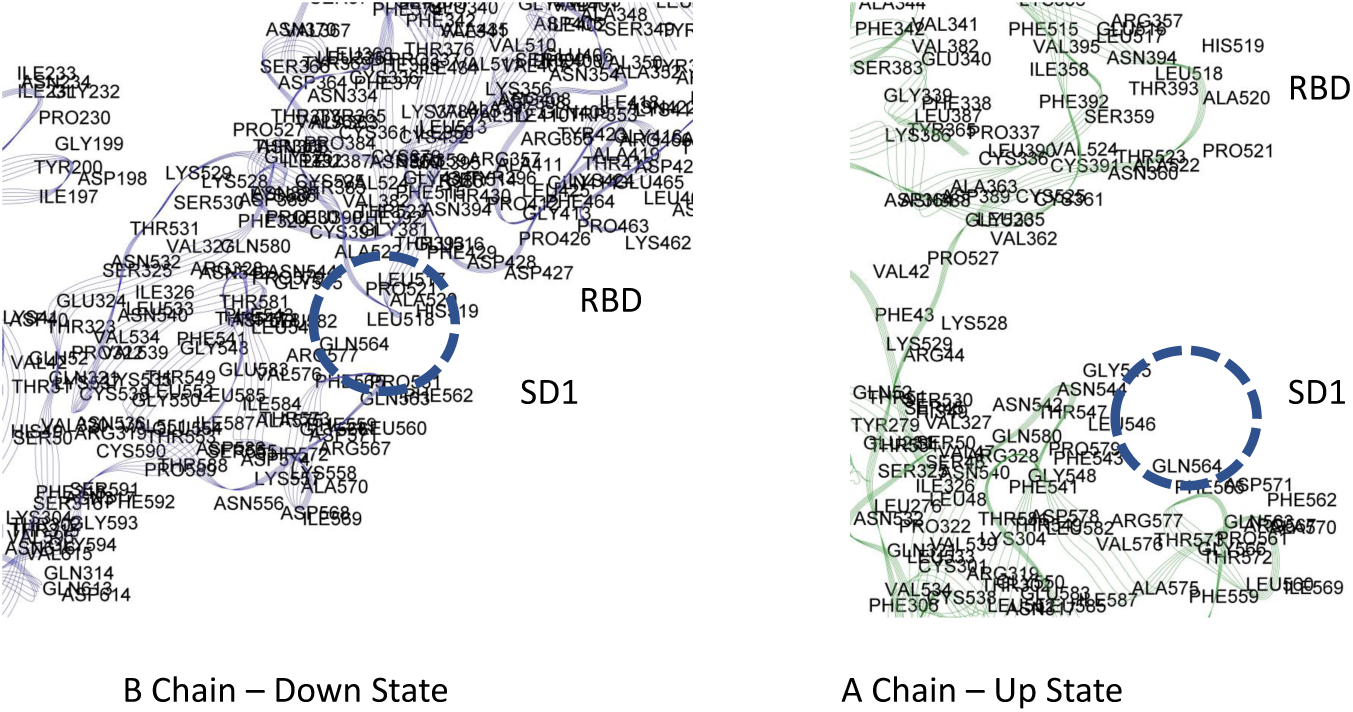
Dominant Energy Landscape for the partial charge interactions between SD1 and RBD of of Chains A and B showing a possible latch for the B Chain.

**Figure 9:**
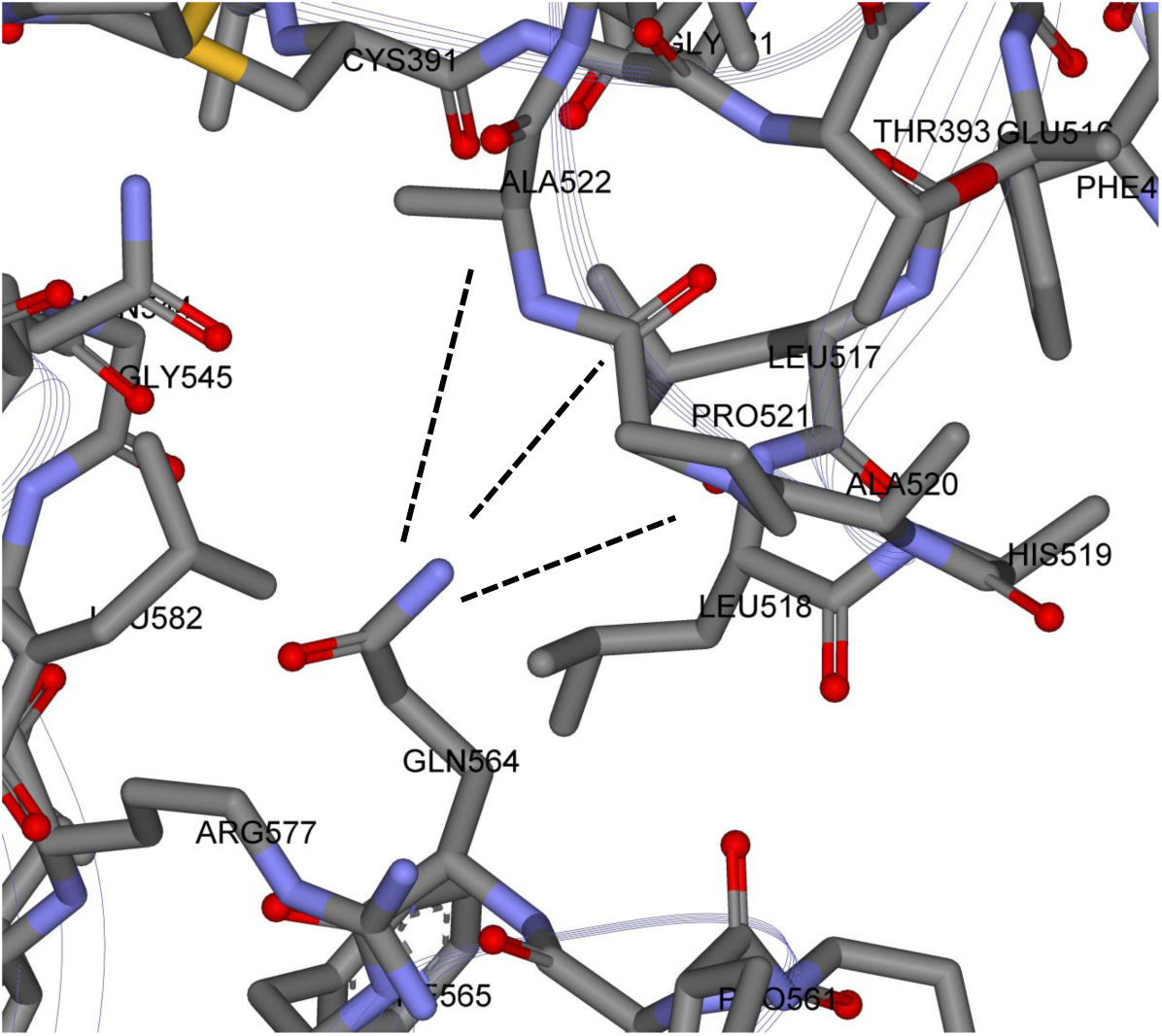
Zoomed View of possible molecular latch between the RBD and SD1 domain: ALA520 PRO 521 ALA 522 with GLN 564.

Interestingly, these are exactly the same three residues of the Up-RBD that participate in the set of dominant energetic interactions with the Down-NTD of its neighbor protomer along the Up-Down interface (Fig.3 previously). Thus, these three dominant energy residues seem to play different roles depending on the Up or Down state of the protomer, which was somewhat surprising.

### Initial Dynamic Test of Latch

In order to determine if these strong, but few residue interactions could possibly function as a molecular latch for the Down state protomer, we conducted some preliminary all-atom molecular dynamics simulation using NAMD2 over a total period of 0.1 *µ* sec. for both a wild S1 chain and a single point mutant S1 chain, both in the same initial down protomer state from PDB ID 6VSB. We used the Charmm-Gui (9) to insert all missing residues from the S1 domain from 6vsb.pdb structure file. For the mutant (single point mutation), we eliminated the R group glue point atoms of GLN 564 by replacing it with GLY 564 and did not alter in any way the three residue sequence ALA520-PRO521-ALA522 in the RBD domain or any other residues throughout the S1 chain. Since our specific interest here was in structure change, we chose not to use alanine screening, which is often used for biological activity mutation analysis to preserve secondary structure. Our trajectory analysis revealed the early release of the RBD from SD1 (within a few nanoseconds) and overall hinge opening and distal release of the RBD for this single point mutant, whereas the hinge angle and RBD relative positions were preserved in the wild state (Fig. 10). (Supplementary Movie: 6VSB-WT vs MT.mov). For completeness, we also determined RMSF values for both wild and mutant (Table S7.latchB.xlxs that clearly showed the relatively high degree of flexibility for the S1 Domain (RMSF values from 5 to 25 Å).

**Figure 10:**
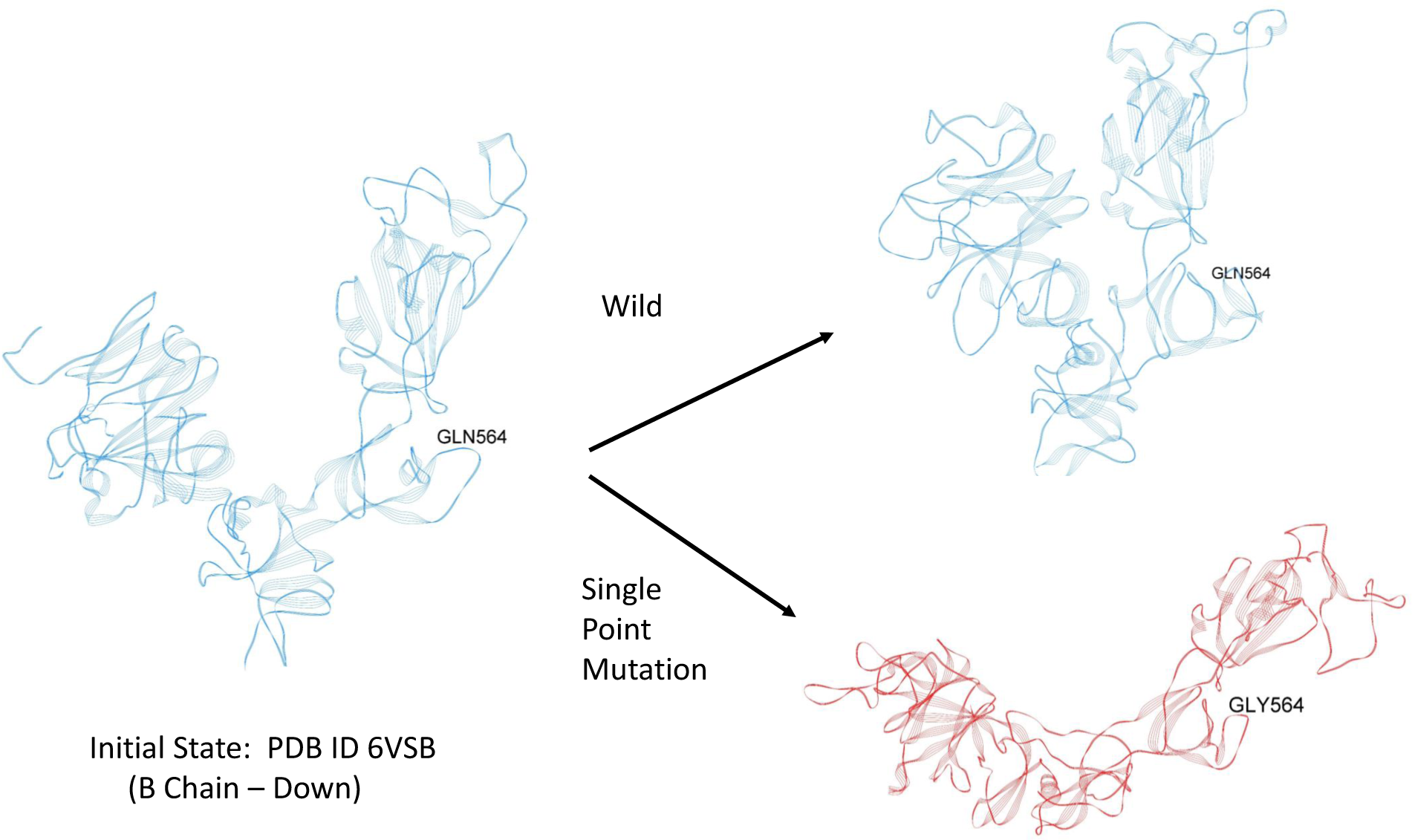
Initial dynamic test of possible latch over a period of approximatley 0.04 *µ* sec. Blue - Wild; Red-Mutant.

These dynamic results are preliminary and more detailed studies, including experimental mutational analysis are needed to further examine the Up versus Down protomer states, including the full trimer state. As we have seen the trimer state is stabilized by cross chain interactions and these may play a critical role in the overall conformation of the complex and the Up versus Down protomer states.

## 4. Discussion

The all-atom energetic mappings are shown to be useful in attempting to understand the overall functionality of the Spike protein of the novel corona virus. The S2, fusion machinery domain, for example, also acts as a rigid base for the more flexible S1, prefusion domain, where flexibility is an important part of its function. The RBD is tethered to the SD1/SD2 domain by two peptide strings: one from the NTD (approximately residues 327-335) and the other connected to the SD1 domains (approximately residues 527-535). These peptide “strings” are stabilized or “tied together” by beta strand structures near the S1/S2 interface. The RBD domain in the Down state protomer is partially stabilized by the dominant energetic interactions of the backbone atoms of three of its residues (in sequence) with the R-group of a single residue from the SD1 domain, which we have called a possible latch. Ironically, these same three residues are involved in helping to stabilize the Up state protomer by interactions with the NTD of its Down state neighbor. This study raises a number of new questions about the origins of the Up and Down state protomers, including the reasons behind the occurrence of the Up state protomer, since as we have shown here, the Down state is partially stabilized by not only the potential latch, but also neighboring chain interactions between the RBD and it’s NTD neighbor. It seems reasonable, however, that the process of “Down to Up” is irreversible in the absence of deliberate outside interference. We further note that the possible three residue latch of the RBD Down state (ALA520 PRO521 ALA522) along with GLN654 of SD1 are conserved across reported variants for SARS-Cov-2 (1). All latch residues are also present in the closely related virions: SARS-Cov-1 and the bat corona virus RatG13; the latter being 96 % homologous to SARS-Cov-2 (1). Preliminary all atom molecular dynamic studies with an in-silico single point mutation of GLN564GLY demonstrated the dramatic structural change of the S1 domain with hinge opening and distal release of the RBD. These preliminary dynamic studies raise a large number of important questions, however, about the long time stability of the Up and Down state protomers and the role of “latch” molecules, exactly how these states originate, the actual transitions that can take place, and if these transitions/instabilities are related in any way to the observed biological and epidemiological behavior of this virus. Nonetheless, our static dominant energy landscape mappings and preliminary dynamic studies will hopefully aid in further understanding of structure-function relationship of the SARS-Cov-2 Spike protein.

## Supporting information

6VSB-WT vs MT.mov

Fig. Suppl.1.jpg

Fig. Suppl.2.jpg

Table Suppl.1.txt

Table Suppl.2.txt

Table Suppl.3.txt

Table Suppl.4.txt

Table Suppl.5.txt

Table Suppl.6.txt

Table.S7.latchB.xlxs

## 5. Acknowledgments

M.H.P. would like to thank the Office of the Vice President for Research, the Department of Chemical and Life Science Engineering, and the Center for High Performance Computing at Virginia Commonwealth University for partial support of our overall research efforts surrounding Covid-19. O.B. was supported by the National Institutes of Health Institutional Research and Academic Career Development Award K12GM119955. M.H.P and O.B. would like to acknowledge the support provided via the COVID-19 HPC Consortium (https://covid19-hpc-consortium.org/), which is a unique private-public effort to bring together government, industry, and academic leaders who are volunteering free compute time and resources, in support of COVID-19 research. The authors would also like to thank all of those involved with this project at the NASA Advanced Supercomputing (NAS) Division, Ames Research Center.

## Open Source Software

OpenContact is freely available under the Third-Party Software Tools listings of the Protein Data Bank: https://www.rcsb.org

## Supplementary Information

**Fig. Suppl.1.jpg**

**Fig. Suppl.2.jpg**

**Tables Suppl.1 - 6.txt**

**Supplementary Movie: 6VSB-WT vs MT.mov**

**Table Suppl.4.xlxs**

## Notes

### Competing Interest Statement

The authors have declared no competing interest.

### Summary of Updates

New long time molecular dynamics results of latch opening via a single point mutation of the Down state protomer.

