## Supplementary figures and images for "Static All-Atom Energetic Mappings of the SARS-Cov-2 Spike Protein with Potential Latch Identification of the Down State Protomer"

### Fig. Suppl.1.jpg

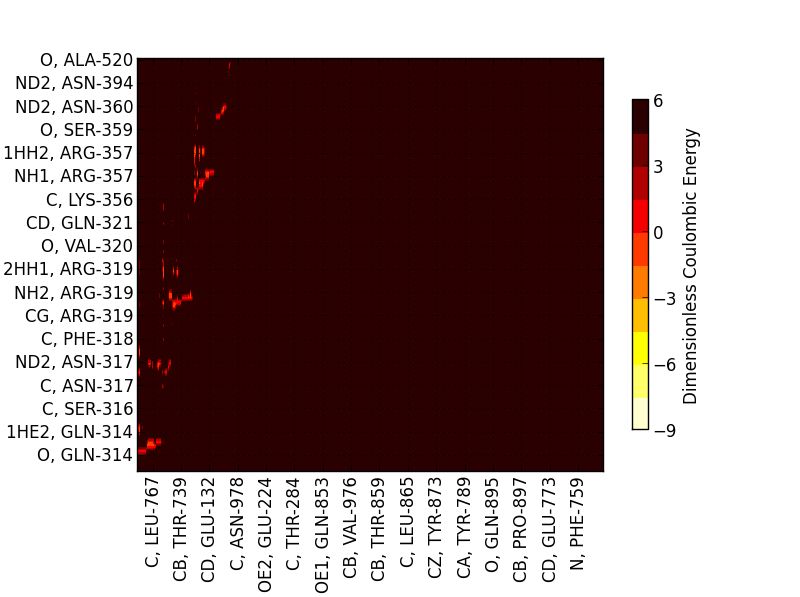

### Fig. Suppl.2.jpg

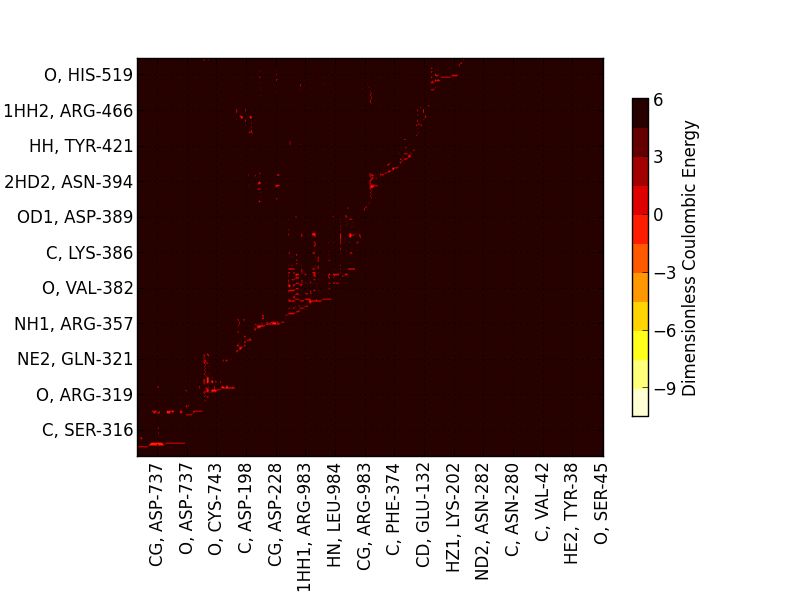
